# Rate versus synchrony codes for cerebellar control of motor behavior

**DOI:** 10.1101/2023.02.17.529019

**Authors:** David J. Herzfeld, Mati Joshua, Stephen G. Lisberger

## Abstract

Control of movement requires the coordination of multiple brain areas, each containing populations of neurons that receive inputs, process these inputs via recurrent dynamics, and then relay the processed information to downstream populations. Information transmission between neural populations could occur through either coordinated changes in firing rates or the precise transmission of spike timing. We investigate the nature of the code for transmission of signals to downstream areas from a part of the cerebellar cortex that is crucial for the accurate execution of a quantifiable motor behavior. Simultaneous recordings from Purkinje cell pairs in the cerebellar flocculus of rhesus macaques revealed how these cells coordinate their activity to drive smooth pursuit eye movements. Purkinje cells show millisecond-scale coordination of spikes (synchrony), but the level of synchrony is small and likely insufficient to impact the firing of downstream neurons in the vestibular nucleus. Further, analysis of previous metrics for assaying Purkinje cell synchrony demonstrates that these metrics conflate changes in firing rate and neuron-neuron covariance. We conclude that the output of the cerebellar cortex uses primarily a rate code rather than synchrony code to drive activity of downstream neurons and thus control motor behavior.

**Impact statement:** Information transmission in the brain can occur via changes in firing rate or via the precise timing of spikes. Simultaneous recordings from pairs of Purkinje cells in the floccular complex reveals that information transmission out of the cerebellar cortex relies almost exclusively on changes in firing rates rather than millisecond-scale coordination of spike timing across the Purkinje cell population.

## Introduction

Normal brain function requires high-fidelity transmission of information from each area to downstream populations of neurons. Therefore, one way to understand how a given brain area influences behavior is to decipher the neural ‘codes’ that relay information to downstream areas. In many cases, information transmission between brain areas occurs via coordinated changes in firing rate^1^ that, passed through synapses, ultimately affect the rate responses of downstream neurons^2,3^. Yet, an alternative exists. Multiple brain areas transmit information via the precise, millisecond timing of spike events^4–6^. Transmission via spike timing requires either a degree of synchrony across the presynaptic population or sufficiently strong synaptic coupling between the upstream and downstream neurons to allow precise temporal information to be deciphered downstream. Rate and temporal information transmission can potentially act in tandem, allowing the simultaneous encoding of multiple stimulus or behavioral features^7–9^.

In many ways, the cerebellum is an optimal structure for identifying the role of synchrony versus rate-based codes for information transmission. First, the ubiquity of the “crystalline” circuit across cerebellar regions has led to the hypothesis that the cerebellar cortex may perform a universal computation^10,11^. Embedded in the universal computation hypothesis is the implicit belief that the codes used for information transmission are conserved across cerebellar regions. Therefore, deciphering the code for information transmission in a single cerebellar location may generalize broadly to other cerebellar regions and tasks. Second, the unique response properties of the sole output of the cerebellar cortex, Purkinje cells (PCs), allow identification of these neurons *in vivo* from extra-cellular recordings alone. PCs fire both high frequency simple spikes (SSs), caused by their intrinsic activity and many parallel fiber inputs, as well as infrequent complex spikes (CSs), caused by the activity of climbing fiber inputs from the inferior olive^12^. Identification of PCs from their unique action potentials allows characterization of the neuronal codes of information transmission in a very specific anatomical pathway: from PCs to their downstream target neurons in the cerebellar nuclei. Third, synchrony has become a plausible neural code for cerebellar output because PC target neurons in the cerebellar nuclei possess unique biophysical properties that may allow precise temporal information to be relayed out of the cerebellar cortex^13^. In particular, the high intrinsic firing rates of cerebellar nucleus neurons^14,15^ and rapid synaptic currents induced by PC firing^13^ could facilitate entrainment of neurons in the cerebellar nuclei specifically following the synchronous spiking of upstream PCs^13,16,17^.

We directly assay the relative contributions of rate and synchrony codes on the output of the cerebellum by taking advantage of (1) a behavior that is readily quantifiable, (2) a cerebellar region where PC output is crucially related to the chosen behavior, (3) the ability to quantify synchrony between simultaneously recorded PCs, and (4) an approach to identify target neurons in the downstream cerebellar nuclei and subsequently quantify the effect of rate versus synchronous PC spiking on their activity. Therefore, we can go beyond previous studies that have characterized PC synchrony in reduced, non-behaving, or even behaving preparations^18–27^, to characterize the effect of PC synchrony signals during a relevant behavior on the responses of identified downstream cerebellar nucleus neurons.

Smooth pursuit eye movements our behavioral paradigm of choice, are readily quantifiable and rely crucially on the floccular complex of the cerebellar cortex^28^. Floccular PCs encode eye kinematic signals^29–33^ where approximately 40-50 PCs^13^ relay this information to each monosynaptically coupled floccular target neuron (FTNs) in the vestibular nucleus^31^. By combining multi-contact electrode recordings in the cerebellar cortex to assay directly PC rate and synchrony codes during pursuit with prior recordings from identified FTNs in the vestibular nucleus^31,34^, we can quantify the codes of cerebellar information transmission in one well-defined neural system. We find that synchrony is, at best, a minor feature of the output code from the cerebellar cortex in this region.

## Results

Our goal was to understand the neural codes that transmit information from the cerebellar cortex to its downstream targets. Here, we assemble data from two connected brain regions to identify the contributions of neuron firing rate and temporal synchrony for the control of a well-characterized motor behavior. We used silicon probes to record the extra-cellular spikes from multiple Purkinje cells (PCs) simultaneously, allowing us to assess the level of spike timing synchrony present in the PC population. Using previous single-electrode recordings from the definitively identified target neurons of the PC population, we quantified the relative contributions of rate and synchrony information relayed to downstream areas. We asked (1) if the spikes of nearby PCs coordinate with millisecond precision to deliver temporally synchronous spiking to downstream areas, (2) whether spike timing synchrony varies over the course of motor behavior, and (3) whether the non-synchronous rate codes of PC firing are sufficient to explain the activity of their target neurons.

### Synchrony between, pairs of Purkinje cells

In the floccular complex of the cerebellar cortex, we recorded *n=32* pairs of well-isolated Purkinje where both PCs were classified unambiguously via the simultaneous recording of their traditional high frequency simple spikes and infrequent complex spikes. The sample of *32* pairs were recorded from *n=44* unique PCs, as we were able to isolate more than a single pair of PCs in several recording sessions. For the example PC pair shown in Figure 1A-D, complex spikes for each unit were easily identifiable and were followed by a stereotypical pause in simple spikes (Figure 1A, B). Based on the contacts of the silicon probe where each PC’s simple spike waveform was the largest (its primary contact), we estimate that these PCs were approximately 200 μm apart. They were in the same Purkinje cell layer and the spikes on each PC’s primary contact were linked to small deflections in the voltage on the other’s primary contact.

**Figure 1.**
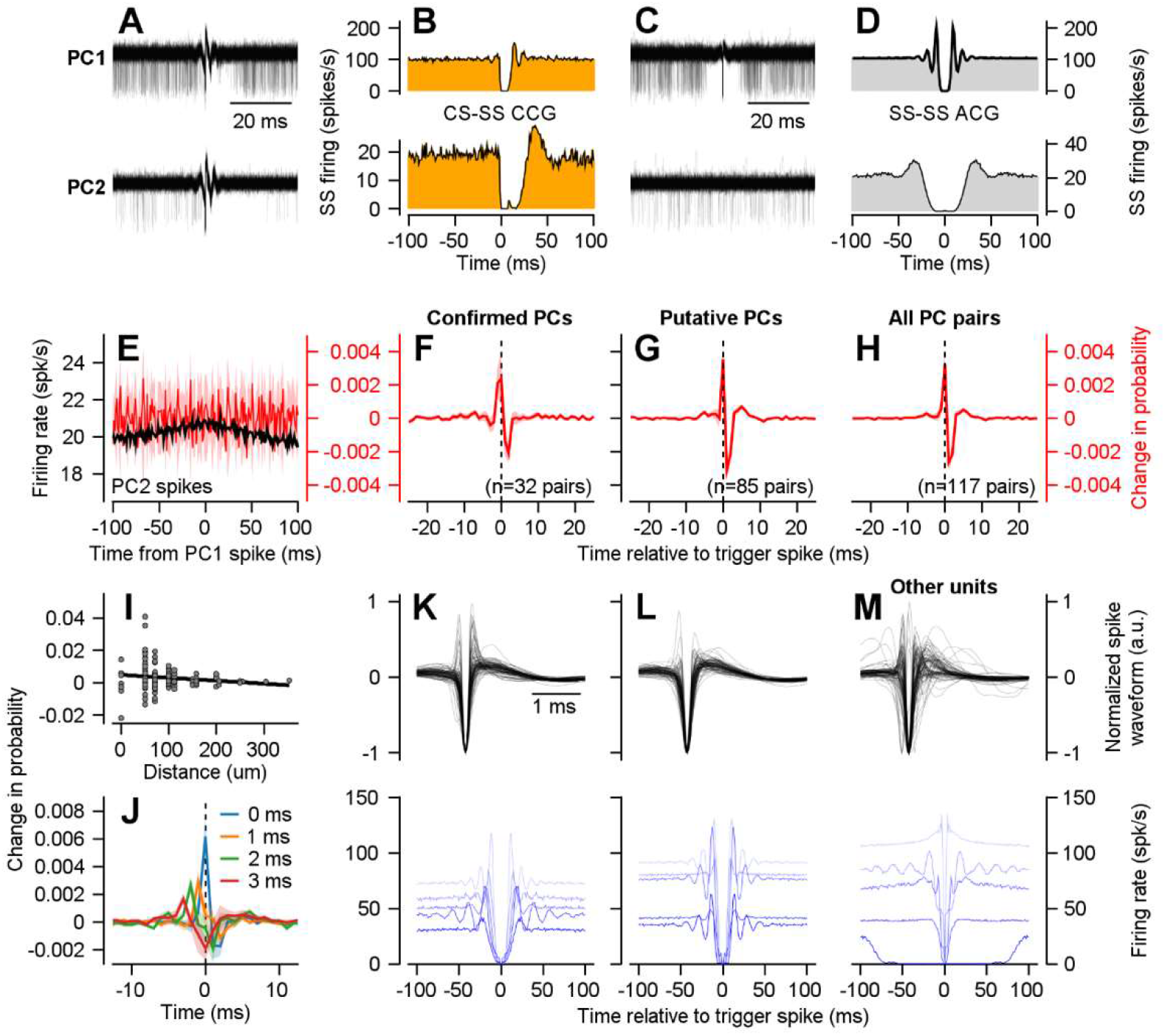
Simultaneously recorded Purkinje cells show small but non-zero spike timing synchrony. **A.** Superimposed raw voltage traces from two PCs, aligned to the onset of a complex spike in each cell. 100 voltage traces are shown for each PC. **B.** Cross-correlograms showing each PC’s simple spikes triggered on the occurrence of a complex spike at *t=0* (i.e., CS-SS crosscorrelogram). **C.** Superimposed raw voltage traces from the same pair of PCs as in (A), aligned to 100 randomly selected PC simple spikes from PC1. **D.** Auto-correlograms triggered on the time of a simple spike for each neuron shown in (A). **E.** Firing of PC2 aligned to the time of PC1’s simple spikes at *t=0* (black). Red curve shows the rate-corrected probability of PC2 firing in 1-ms bins in units of change in probability (right axis). Shaded region denotes 95% confidence intervals. **F-H.** Rate-corrected probability of firing averaged across a population of simultaneously recorded PCs (**F**), putative PCs that lack a recorded complex spike (**G**), and across all PC and putative PC pairs (**H**). Shaded regions denote SEM across PC pairs. **I.** Rate-corrected probability of synchrony versus distance between the primary contact for pairs of simultaneously recorded PCs. Black line denotes the best linear fit. **J.** Rate-corrected CCGs separated according to where the maximum value occurred between the *t=0* to *t=3* millisecond bins. **K-M.** Primary channel waveform (top) and example auto-correlograms (bottom) for populations of known PCs (**K**), expert-identified putative PCs (**L**), and randomly selected non-PCs (**M**).

Visual inspection of PC2’s spike-triggered responses demonstrated little discernable change in the occurrence of PC2’s simple spikes due to the firing of PC1 (Figure 1C, bottom trace). The absence of strong synchrony is confirmed by the cross-correlogram (CCG) of the simple spikes of PC2 triggered on those of PC1 (Figure 1E, black trace). We constructed the CCG as:

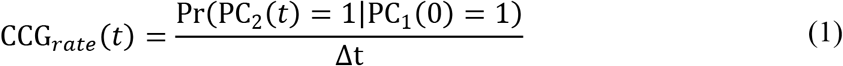

In words, Equation (1) describes the probability of observing a spike in PC2 at time *t* relative to a spike of PC1 at *t*=0. The normalization factor, Δ*t*, ensures that the magnitude of CCG is not dependent on the chosen temporal resolution (1 ms in Figure 1E) and yields a normalized CCG with units of spikes/s. The CCG between PC1 and PC2 in Figure 1E shows a small, broad, increase in the firing rate of PC2, almost certainly because of slow co-modulation of the PC firing rates due to behavior-related firing. We isolated the true synchrony in simultaneously record neurons by removing the co-occurrence of spikes expected solely by changes in either of the neurons’ firing rates (see Methods), resulting in a “rate-corrected CCG” (red trace, Figure 1E). The corrected CCG does not deviate from the 95% confidence intervals expressed in either probability or rate, indicating an absence of millisecond scale synchrony in these 2 PCs.

Across our full sample of PC-PC pairs, we observed low levels of spike timing synchrony. There was a small narrow peak near time zero in the mean CCG (Figure 1F) for the 32 pairs of PCs identified definitively by the post-complex spike pause in simple spike firing. The occurrence of a spike in the trigger PC resulted in an increased probability of observing an unexpected synchronous spike of 0.0024 ± 0.0016 (mean ± SEM) in the second unit, a difference that was not significant across the population (one sample t-test, t(31)=1.48, p=0.15). Note that we determined which neuron of a PC-PC pair was the “trigger” neuron based on the shape of the rate-corrected cross-correlogram (see Methods). As the mean simple spike firing rate of the 44 PCs in this sample was 47.7 ± 3.6 spikes/s, the probability of synchrony we observed would result in an extra 0.11 spikes/s (47.7 x 0.0024) when the two PCs fire together, beyond the 2.3 synchronous spikes/s that would be expected due to random chance. The peak in the CCG at time zero was very similar (Figure 1G) when we included pairs where one or both of the pair were identified as putative PCs by their laminar location, waveform, and resting discharge properties, without the presence of a clear climbing fiber response (see below); the synchrony was statistically significant but still small (an excess probability of 0.0035 ± 0.0009 synchronous spikes when the trigger neuron fired) for the population of putative PCs (t(84)=3.88, p<0.001).

For the full population of 117 PC-PC pairs from known PCs plus putative PCs, the firing of the trigger PC resulted in an excess probability of synchronous spikes of 0.0032 ± 0.0008 in the second PC (t(116)=4.1, p < 0.001). We found no significant difference in synchrony between our sample of known-known and putative-putative PC pairs (independent samples t-test, t(115)=-0.60, p=0.52), nor for comparison of pairs of known PCs to a non-overlapping sampling of *n=49* known-putative PC pairs (independent samples t-test, t(79)=-0.83, p=0.41). Across the 133 unique neurons in this sample, the mean simple spike firing rate was 59.6 ± 2.3 spikes/s, suggesting that the excess synchrony we measured would result in an extra 0.19 spikes/s when the two PCs fire together, beyond the 3.6 spikes/s expected by chance.

Several features of our data make us think that the near-zero synchrony we record is real and is not an artifact of errors in spike-sorting. (1) We curated our spike-sorting carefully and only included neurons with a high signal-to-noise ratio (Figure 1A, C). (2) We tested our spike-sorting pipeline with artificial data that mimicked our use case and verified the veracity of the CCGs produced under simulated real-world conditions^35^. (3) We accepted only PCs with autocorrelograms (ACGs) that showed essentially no evidence of refractory period violations (Figure 1D). (4) The non-zero CCGs for our PC-PC pairs showed a peak followed by a trough of approximately equal integrated magnitude (Figure 1F-H), a feature that would not be expected in CCGs if millisecond-scale synchrony is an artifact of spike-sorting error (see Discussion).

Our analysis of spike timing synchrony between PCs revealed two additional important features. First, the majority of our non-zero values of synchrony came from pairs of PCs that were quite close together (Figure 1I), including some recorded on the same primary contact. We found a significant effect of distance on the magnitude of the synchrony between pairs (Spearman ρ=-0.19, p=0.04). In addition, the variance of measured synchrony significantly decreased with distance (Spearman ρ=-0.94, p < 10^-4^). Here, our proxy for distance was based on the electrode contact with the largest simple spike waveform for each PC in the recorded pair. The 50 μm separation between contacts on our probes limited the resolution of distance, e.g.: 0 μm (same contact), 50 μm (adjacent contacts), 71 um (diagonal contacts). We found essentially negligible levels of synchrony for pairs that were separated by more than 100 μm (probability of excess spikes of 0.0011 ± 0.0005, mean ± SEM, given a spike in the trigger neuron). PCs separated by less than 100 um showed somewhat larger levels of synchrony (excess probability of 0.0041 ± 0.001 when the trigger neuron fired, equivalent to an excess 0.24 synchronous spikes/s). Second, we observed a broad distribution of the timing of the peak in the CCGs for different PC-PC pairs. In our population of *n=117* PC pairs, 47 pairs (40.2%) showed a peak within 1 ms of each other, suggesting that the other half of the pairs featured either no peak or a peak in the CCG that did not correspond to millisecond level synchrony. Consider, for instance, the 22 PCs summarized by the red curve of Figure 1J. They showed a peak in their firing 3 ms before the trigger neuron fired and, at the time of the trigger neuron’s spike, a decrease in our measure of synchrony due to the mandatory trough following the peak at t=-3ms.

We validated the inclusion of putative PCs in our analysis by applying several criteria. Overlaying the extra-cellular waveforms from our population of known PCs (Figure 1K, top) demonstrated consistent across-PC temporal features. Similarly, confirmed PCs showed stereotypical ACGs featuring multiple lobes with resting discharge rates between 30 and 100 spikes/s (Figure 1K, bottom). Therefore, we manually classified a neuron recorded without an associated complex spike as a putative PC based on (1) its location in the Purkinje cell layer, (2) its extra-cellular waveform, (3) its auto-correlogram. Neurons that we classified as putative PCs showed the same features as confirmed PCs (Figure 1L), whereas an equal number of randomly selected “other” units recorded in the granule or molecular layers showed a wider range of waveforms and ACG shapes (Figure 1M).

Taken together, our results demonstrate that pairs of PCs show coordinated timing of simple spikes with a magnitude that is above the level expected solely from their joint firing rates. However, millisecond spike timing synchrony across the population was small and variable, even for pairs of PCs that were very close together.

### Coincident firing of PCs that share the same complex spike field

One theory suggests that Purkinje cells that share a climbing fiber input might show higher-levels of spike timing synchrony^23,25^ and project to a common target neuron^36–38^, creating a local synchrony signal that would focus on individual target neurons in the cerebellar nuclei. Approximately 10 local PCs share the same climbing fiber input^39^, and events on subsets of climbing fibers can be synchronized^40,41^. If synchronous complex spikes can drive simple spike synchrony, and PCs that share a climbing fiber input project to same neuron in the cerebellar nucleus, then we might expect substantially more synchrony that would be meaningful for downstream signaling from PC pairs that share the same complex spike response.

The theory of simple-spike synchrony tied to climbing fiber synchrony is not supported by our recordings from a population of *n=5* confirmed PC pairs that appeared to receive inputs from the same or highly-synchronized climbing fiber inputs. The evidence for common climbing fiber inputs comes from the finding that both PCs in simultaneously recorded pair paused for the same complex spike (Figure 2A). We found limited simple-spike timing synchrony, comparable to what we found in the full sample of PC-PC pairs, in the CCGs for both the example pair of PCs in Figure 2A and the full population of 5 pairs of PCs (Figure 2B, excess probability of 0.0046 ± 0.0059 when the trigger neuron fired, t(4)=0.78, p=0.48). We conclude that synchronous climbing fiber inputs probably do not preferentially synchronize the simple spike activity of the PCs they contact. Thus, neither shared CS inputs nor location within a given cerebellar microzone is sufficient to create more than a tiny amount of millisecond synchrony among the simple-spike responses of neighboring PCs.

**Figure 2.**
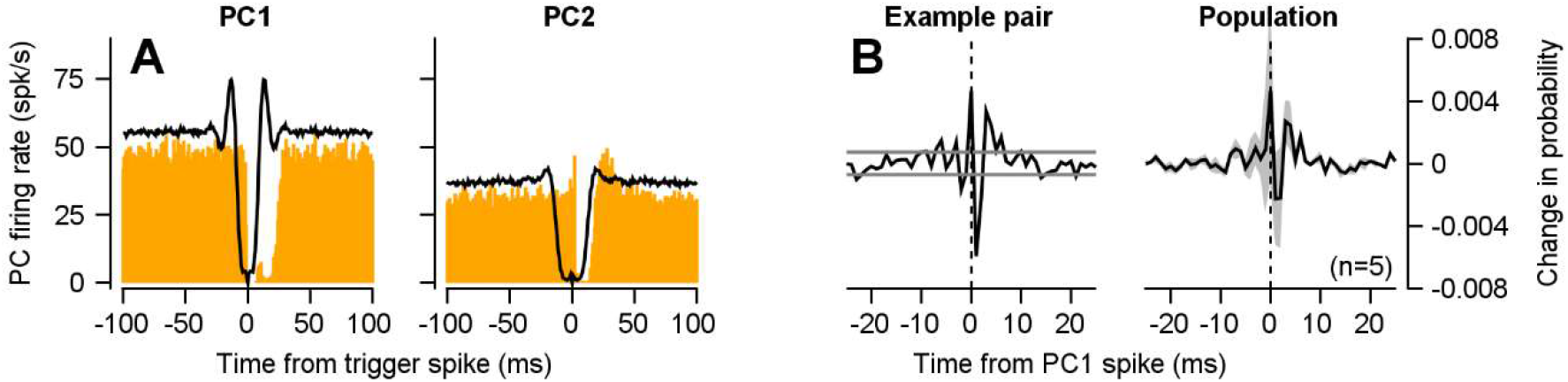
Synchrony among Purkinje cells that share the same complex spike response mirror the complete population. **A.** Example ACGs (black) and CS-SS CCGs (orange) for two Purkinje cells that pause their simple spike responses following the same complex spike. **B.** Rate-corrected CCGs for the example pair shown in (**B**) (left) as well as across a population of five PC pairs that show pauses to the same complex spike (right). Shaded regions in (**B**) denote SEM across the five PC pairs.

### Lack of synchrony modulation during movement

Our analysis to this point measured synchrony across complete recording sessions and thus may hide strong synchrony that exists specifically during movement or in specific phases of a movement. As the floccular complex of the cerebellum is crucial for the execution of smooth pursuit eye movements^28,42^ and floccular PCs respond robustly during pursuit^29,32,33,43^, we asked whether pairs of floccular PCs preferentially synchronize or desynchronize their activity at any specific moments during execution of pursuit.

Purkinje cells show direction-selective modulation of simple-spike firing during pursuit, but we found no evidence for modulation of synchrony during movement. The example PC in Figure 3A showed increased firing during pursuit toward the side of the recording, defining its preferred or “SS-on” direction, and decreased firing for pursuit in the opposite direction (“SS-off”). In almost all cases, the PCs in our simultaneous recordings shared the same SS-on and SS-off directions (Figure 3C), favoring pursuit either in the ipsiversive direction (toward the side of recording, 52%) or downwards (30%) (Figure 3B). As others have found, the CS-on direction in our sample was usually opposite the PC’s SS-on direction (see Figure 3A); for most PCs, the CS-on direction was contraversive (Figure 3B) and the CS-on directions were shared by the PCs in most pairs (Figure 3C). The angular difference between the SS-on directions of simultaneously recorded PCs had circular means of 14.7° (red vertical line) that was not significantly different than zero (t(116)=0.69, p=0.17). This distribution was non-uniform as suggested by a circular dispersion of 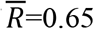 (the circular dispersion is zero when angles are uniformly distributed and one when representative of a single angle). The CS-on directions between simultaneous pairs were also similar: mean angular difference of 6.3° (t(31)=-1.1, p=0.2, circular dispersion of 0.84).

**Figure 3.**
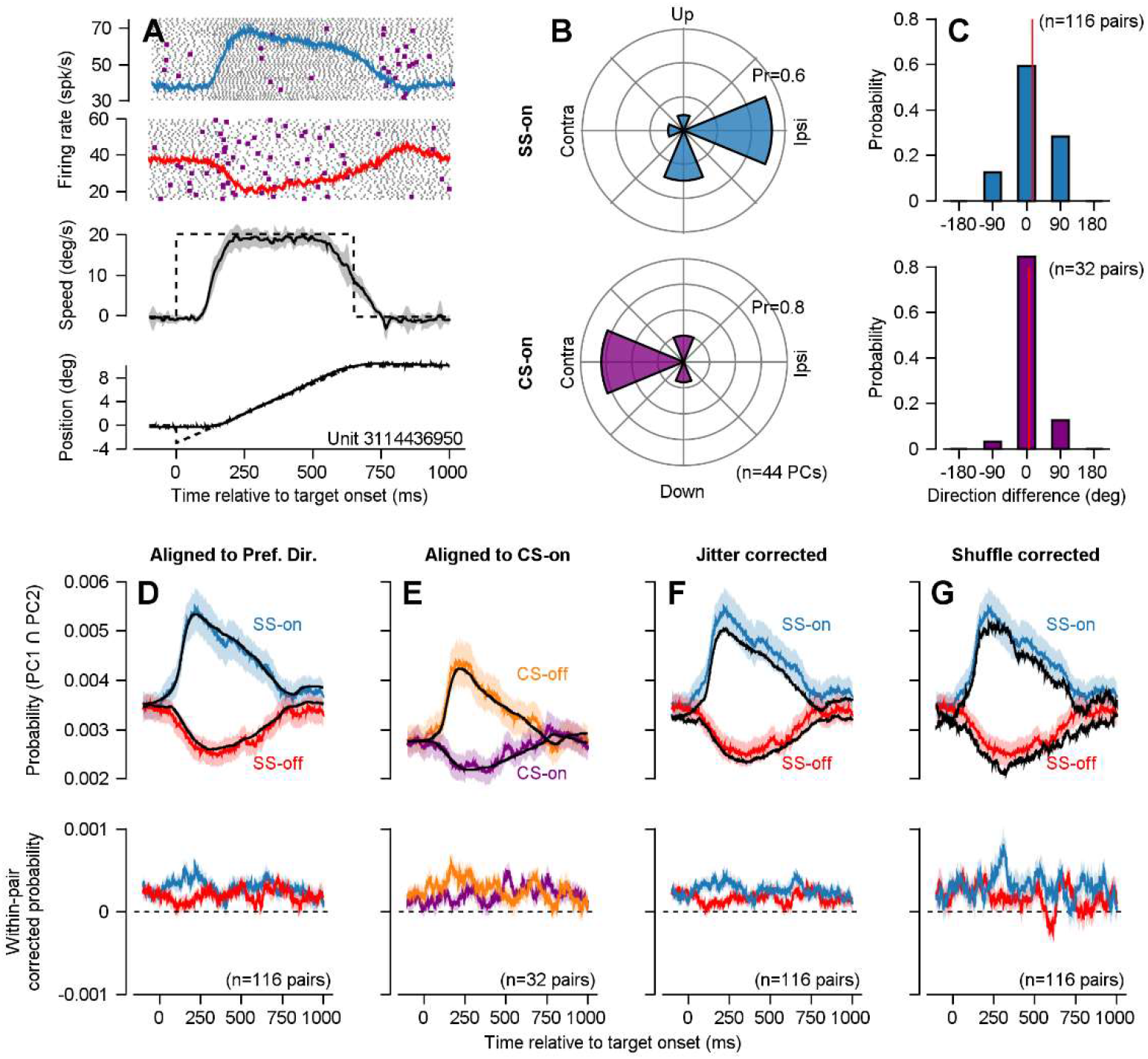
Purkinje cells do not synchronize preferentially at any time during pursuit eye movements. **A.** Firing rates and rasters for an exemplar Purkinje cell during smooth pursuit in the preferred (top raster) and anti-preferred (bottom raster) pursuit directions. Blue and red traces show average firing rate across time. Black and purple dots denote simple and complex spikes, respectively. Example eye (black) and target (dotted) velocity and position traces for a pursuit trial with a pursuit speed of 20 deg/s appear in the bottom panels. Gray shaded regions denote 95% confidence intervals across pursuit trials. **B.** Probability distributions of preferred directions for simple spike (top, blue) and complex spikes (bottom, purple) populations of Purkinje cells. **C.** Pairwise angular difference between preferred simple spike (top, blue) and complex spike (bottom, purple) directions for simultaneously recorded Purkinje cells. Red vertical lines denote the angular mean across all pairs. **D.** Probability of observing millisecond-scale synchrony between pairs of simultaneously recorded Purkinje cells in the preferred simple spike (blue) and anti-preferred (red) directions for 20 deg/s pursuit. Black lines show the rate-corrected null probabilities of intersection at millisecond time scales. Bottom plots show the within-cell measured probability of synchrony minus the rate-corrected null probabilities. **E.** Same as for (**D**), but relative to the preferred CS direction (CS-on). **F.** Same as (**D**) except null hypothesis probabilities are computed using the jitter-corrected method with windows of 5 milliseconds. **G.** Same as in (**D**), except null hypothesis probabilities are computed by shuffle correction across trials. Shaded regions in **D-G** denote SEM across PC pairs.

As expected from the common SS-on directions within PC pairs, there was an increase in the probability of observing coincident spikes from PC2 during pursuit of a 20 deg/s target in PC1’s preferred direction (Figure 3D, blue). For pursuit in the SS-off direction of PC1, both PC1 and PC2 tended to decrease their firing rates together, resulting in a decrease in the probability of observing the cooccurrence of spikes within the same millisecond (Figure 3D, red). Subtraction of the synchrony expected from change in firing rates alone confirms the expectation that the probability of coincident spikes follows the expected effect of firing rate. Here, we jittered the exact timing of spikes in PC1 and PC2 under the assumption of a uniform probability of firing between adjacent interspike intervals (see Methods). Jittering destroyed any temporal information shared in the exact timing of PC1 and PC2’s spikes but retained the mean firing rates of the two neurons within a local window. The product of the jittered spike timeseries of PC1 and PC2 predicts the probability of coincident spikes due to changes in firing rate alone (black traces in Figure 3D). The resulting difference between the observed and rate-only estimated curves across pairs of PCs (the ‘rate-corrected synchrony’, bottom graphs in Figure 3D) is effectively uniform across the pursuit movement, and only slightly positive. Excess synchrony in the SS-on direction was 0.27 ± 0.06 spikes/s (mean ± SEM, one sample t-test, t(115)=4.40, p<10^-3^). Excess synchrony in the SS-off direction was similarly small: 0.18 ± 0.06 spikes/s (t(115)=3.07, p=0.003). Comparison of excess synchrony between the SS-on and SS-off directions showed no significant effect of pursuit direction (paired samples t-test, t(230)=1.01, p=0.31), further suggesting that the baseline level of synchrony we observed is unaffected by motor behavior.

We performed several control analyses. When we aligned the PC responses to the CS-on versus CS-off directions of PC1, the measured synchrony decreased versus increased during the pursuit trial (Figure 3E, purple), as expected given that the CS-on direction is usually the same as the SS-off direction (Figure 3B). Again, the actual synchrony was almost identical to that predicted given the firing rates of the PCs (Figure 3E, black curves) and the rate corrected synchrony (Figure 3E, bottom) was slightly greater than expected given the firing rates of both PCs but was not dependent on pursuit direction or time in the pursuit trial. The same results emerged when we used jitter-correction (see Methods) to remove temporal information with timescales greater than 5 ms (Figure 3F) as well as when we shuffled the data by randomly permuting the order of the pursuit trials for one of the PCs in each pair (shift predictor, Figure 3G).

### Synchrony index

A previous publication reported that spike timing synchrony between pairs of PCs in the oculomotor vermis shows a large increase specifically when the firing rate declines at the end of saccadic eye movements^27^. The conclusion in that paper relies on a metric of synchrony^19,27^ called the “synchrony index”, which quantifies the probability of observing coincident simple spikes across PC pairs divided by the product of their independent probabilities.

To allow direct comparison with previously reported measurements of synchrony, we used the synchrony index to reanalyze the simple spike responses of pairs of PCs over the course of pursuit eye movements in the CS-on and CS-off directions. For an exemplar pair of PCs that showed simultaneous decreases in SS firing during pursuit in their shared CS-on direction (Figure 4A), the synchrony index (Figure 4B, green trace) implies an increase in synchrony at the time firing rate decreases. The same result appears in averages across our full sample of PC-PC pairs (Figure 4C). Yet, the rasters in Figure 4B seem to belie the conclusion of increased synchrony at that time. The red symbols indicate the times of spikes that occurred within 1 ms in the two PCs, and do not suggest any increase in synchrony at the time when the synchrony index shows a peak.

**Figure 4.**
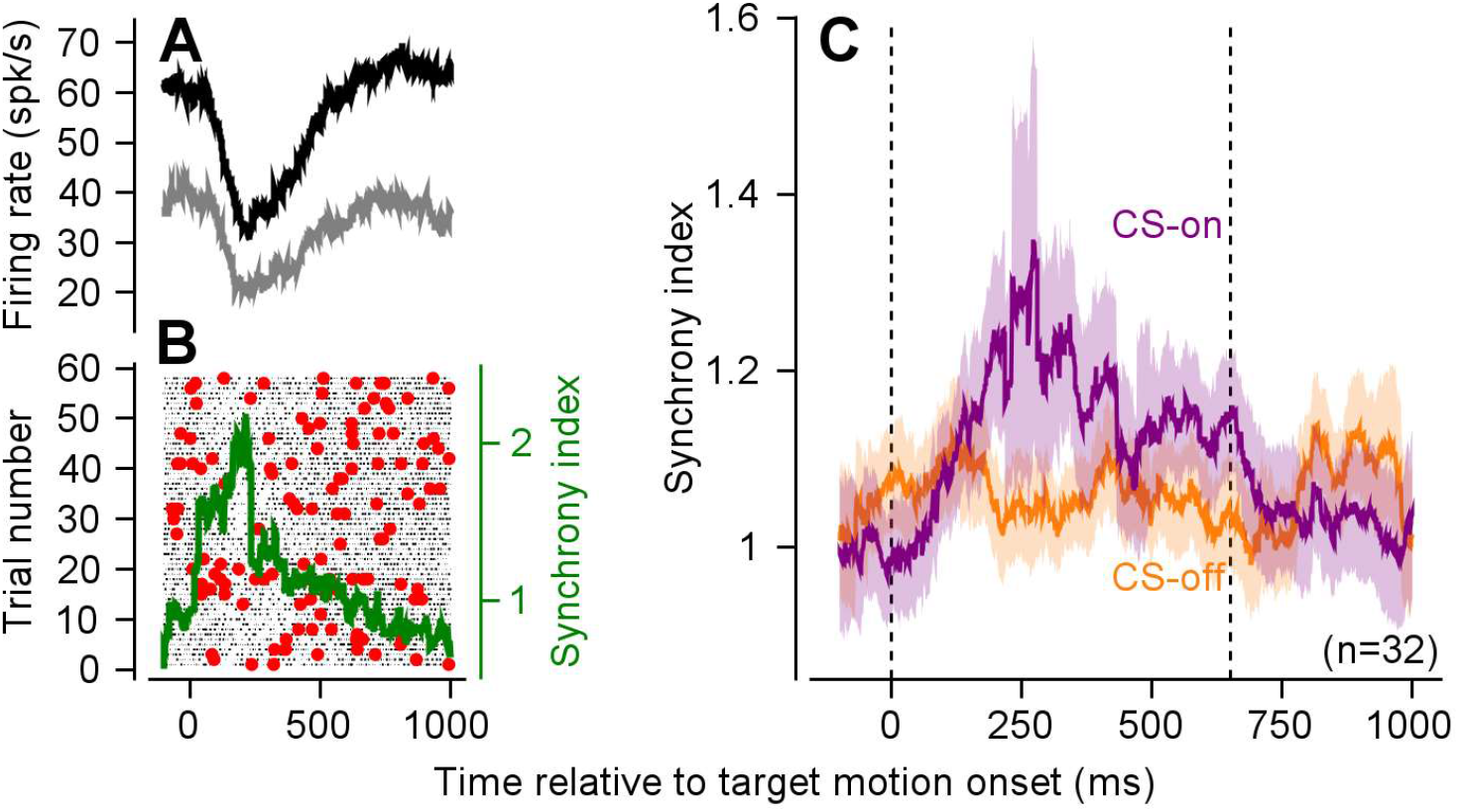
Alternative metrics to assay synchrony incorrectly discover temporally-specific Purkinje cell coordination. **A.** Exemplar paired PC recordings measured during smooth pursuit eye movements in the CS-on direction. The two traces show mean firing rates during pursuit for the two neurons in a pair. **B**: Rasters for each neuron across smooth pursuit trials. Spikes that occurred in the same millisecond bin in the two neurons are plotted as red symbols. The green trace shows the synchrony index averaged across trials. **C.** Synchrony index (± SEM) across *n=32* pairs of PCs in the CS-on (purple) and CS-off directions (orange). Data provide a direct comparison to the bottom graph in Figure 3E.

The discrepancy between our conclusions based on rasters of coincident firing and those suggested by the synchrony index arises because the synchrony index is proportional to both the coherence of spiking of the two PCs and the inverse of the firing rates of the two neurons (see derivation in Methods). Our analysis and simulations based on the firing rates of PCs in the oculomotor vermis (Supplementary Figure 1) confirms that synchrony index misleadingly increases and can become quite large when firing rate decreases, even when we contrive spike timing synchrony to be constant across time.

### The contribution of synchrony to downstream neuron firing

Our final goal is to move from measures of the temporal specificity of spiking across pairs of simultaneously recorded PCs to an assessment of what information (if any) is relayed to downstream neurons in the deep cerebellar nuclei by synchrony in the population of 40-50 PCs that converge on a single downstream unit^13^ (Figure 5A). First, we assessed the synchrony in a simulation of the inputs to a downstream neuron based on the measured distribution of rate-corrected covariance values between pairs of simultaneously measured PC spike trains (Figure 5B-D). Second, we asked whether the firing of downstream neurons recorded previously during pursuit could be accounted for based solely on the firing rates of PCs (Figure 5E-H).

**Figure 5.**
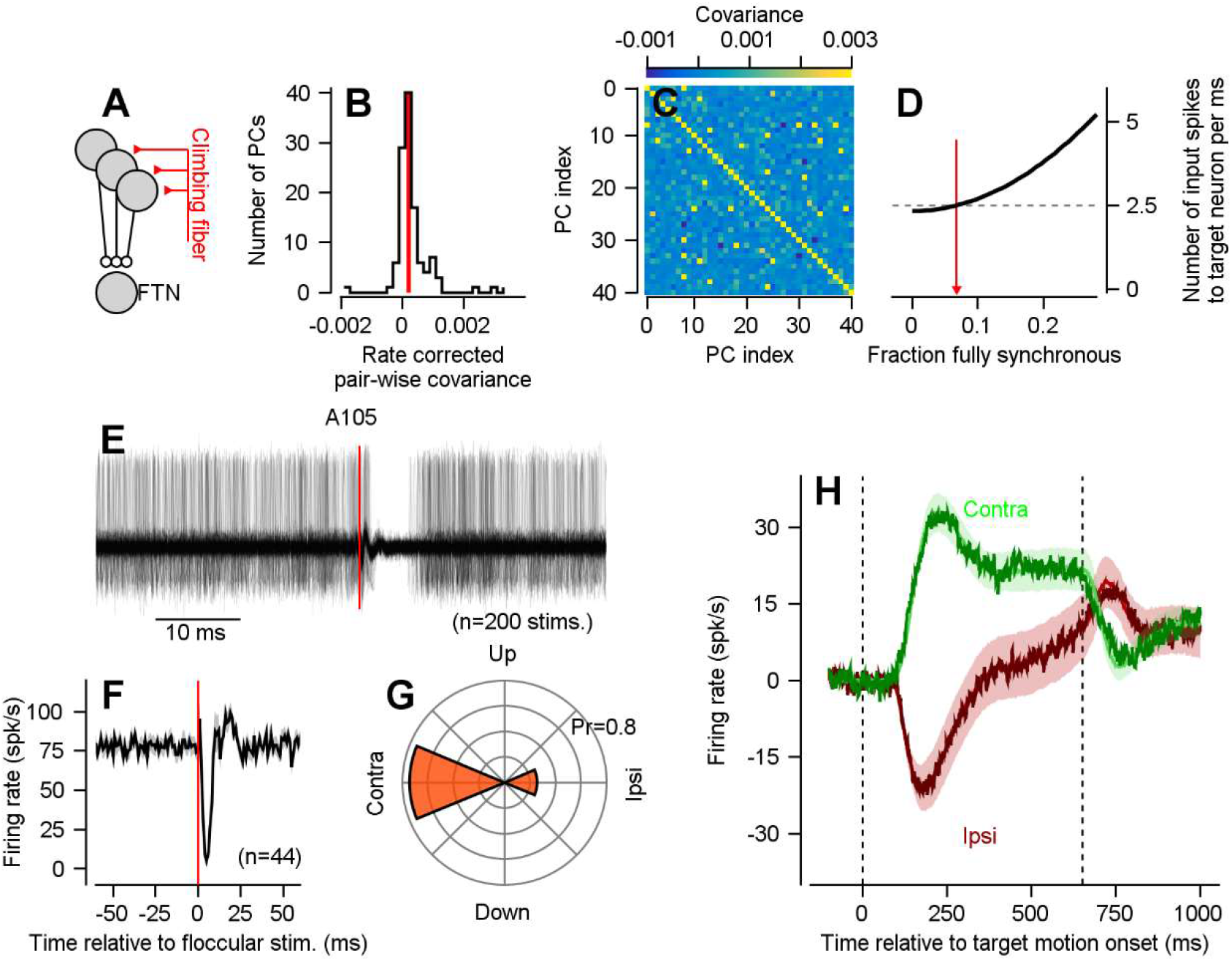
No evidence for a temporal code for transmission of information from Purkinje cells to downstream neurons. **A.** Schematic diagram showing the convergence of a subpopulation of PCs that share the same climbing fiber input onto a single floccular target neuron (FTN) in the vestibular nucleus. **B.** Distribution of rate-corrected covariance values for all PC-PC pairs in our dataset. Red vertical line represents the mean across *n=117* pairs. **C.** Simulated pair-wise covariance matrix for a population of *n=40* model PCs that provide inputs to a single model FTN. **D.** Black curve shows the number of synchronous input spikes to the model target neuron in a given ms as a function of the fraction of input PCs with synchronous spikes. Red arrow denotes estimated fraction of fully synchronous spike trains from the simulated population of neurons with pair-wise covariances shown in (**C**). Panels **E-I** show reanalysis of data from a previous publication^27^. **E.** Exemplar recording from a floccular target neuron that receives monosynaptic inputs from floccular Purkinje cells. Plot shows *n=200* superimposed voltage traces aligned to the onset of single shock stimulation (red vertical line) in the floccular complex. **F.** Average firing rate responses across a population of *n=44* FTNs, aligned to the onset of stimulation. **G.** Probability distribution of preferred directions of smooth pursuit for all FTNs. **H.** Firing rate responses of FTNs for contraversive (green) and ipsiversive (red) pursuit. The pale shading shows the mean + 1 SEM of the measured firing rate. Lines show the best fit predictions from the population of Purkinje cells recorded for this paper.

In our first step, we simulated a population of *n=40* PCs with pairwise covariances drawn from the empirical distribution (Figure 5B, mean rate-corrected covariance: 2.1×10^-4^ ± 5.3×10^-5^) as well as mean firing rates taken from our PC population (59.6 ± 2.3 spikes/s). Viewed from the perspective of a downstream neuron, the measured degree of synchrony in PC-PC pairs predicts that the input stream will be slightly increased to 2.50 ± 0.16 spikes/ms at the time of a spike in one of the 40 input PCs (mean ± SD across 50 bootstrapped PC populations), compared to a uniform input of 2.31 ± 0.18 spikes/ms for completely independent spike trains in the 40 inputs PCs. The mean rate was the same for both the independent and correlated populations, but due to the difference in covariances between the two populations, the distribution of spike timings differed. For comparison with previous literature, we then used the predicted excess in simultaneous spiking to predict what percentage of spike trains would be fully synchronized based on an easily derived relationship (Figure 5D, black curve, see Equation 8). The measured degree of synchrony is equivalent to having identical spike trains in slightly more than a single pair of the 40 PC inputs (1/40, ~5%) and completely independent spikes in all other PCs in the input population. Previous in *vitro* analysis demonstrated that 5-10% synchrony, coupled with baseline firing rates near 60 spikes/s, would result in minimal entrainment of downstream nuclear neurons^13^ (Figure 3C of that report).

Our second step took advantage of the fact that we have recorded both from pairs of PCs and, in separate experiments, from the identified target neurons of the same PCs. In our previous study, we identified floccular target neurons (FTNs) in the vestibular nucleus by implanting a chronic bipolar stimulating electrode in the floccular complex and identifying neurons that showed inhibition (Figure 5E, F) at monosynaptic latencies^31,34^. During pursuit eye movements, FTNs demonstrated preferred directions that were biased for contraversive pursuit (74%, Figure 5G), as expected if they are inhibited by PCs that generally prefer ipsiversive pursuit (Figure 3B).

We could reproduce the firing of FTNs during pursuit in the ipsiversive and contraversive directions with a simple linear model of PC firing rates, without assuming spike timing synchrony in excess of that expected from the firing rates of the PCs. The model was:

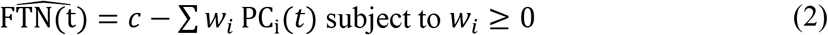

where *w_i_PC_j_*(*t*) represents the non-negative weighted contribution of the *i^th^* PC to the firing rate of the FTN and *c* represents the FTN’s background firing rate. Across *n=39* FTNs, the linear combination of our full sample of PC rate responses was able to account for almost all of the variance in FTN firing (Figure 5H) for both contraversive and ipsiversive pursuit (R^2^=0.96 ± 0.04 and 0.98 ± 0.02, mean ± SD). When we restricted the PC population to 40 randomly selected PCs, we still could account for approximately 90% of the variance of FTN firing (R^2^=0.93 ± 0.006 and 0.89 ± 0.008 for ipsiversive and contraversive pursuit, mean ± SD across 50 bootstrapped PC populations), suggesting there is sufficient variability in the PC population to account for the majority of the variance in FTN responses. Synchrony might account for the fraction of the unexplained variance, but the magnitude of such a contribution seems to be small.

## Discussion

How does the cerebellar cortex relay information to downstream structures? Does it use a traditional rate code where the downstream neurons simply fire in relation to the weighted sum of the post-synaptic potentials in their inputs? Or is millisecond timing of cerebellar output critical in determining how downstream neurons fire? Because of its well-understood anatomy and physiology, the cerebellum seems like an excellent structure to quantitatively address the critical, general question of how a neuron’s output results from its input spike trains. Indeed, the physiology of neurons in the cerebellar nucleus^13–15^ suggests a potential mechanism for signaling by millisecond synchrony in incoming spike trains and previous results by Person and Raman^13^ elegantly demonstrated its plausibility.

### No evidence for Purkinje cell synchrony as a neural code

We found essentially no evidence for millisecond synchrony as a neural code in a specific part of the cerebellum called the floccular complex. The part of the cerebellum we studied is particularly advantageous to address questions of neural information transfer because we understand a great deal about floccular anatomy, physiology, and function. Floccular PCs discharge reliably in relation smooth pursuit eye movements^29,32,33,43^, previous recordings from neurons identified as their primary targets (FTNs) allowed us to assess the effect of PC firing on floccular output neurons^31,34^, and modern multi-contact probe technology allows us to record from many pairs of nearby PCs with single spike temporal and spatial resolution.

Three pieces of evidence convince us that the flocculus does not use synchrony as the primary, or even important mechanism of information transfer to FTNs. First, the objectively measured level of PC synchrony is small. Second, we did not observe a change in the level of synchrony during pursuit eye movements. Third, FTN responses during pursuit could be well approximated by Purkinje cell firing rates.

We also did not find evidence of millisecond synchrony organized according to climbing fiber projections^39,44^, which define cerebellar ‘micro-zones’. Millisecond synchrony of simple spikes responses within a micro-zone was small and similar to the overall population of PC-PC pairs. As a result, we think that the small synchrony we observe in nearby PCs probably results from local circuit properties and mossy fiber inputs rather being driven by climbing fiber inputs^26^.

Person and Raman^13^ showed the synchrony *could be* part of the cerebellar output code. They explained how a code based on a high degree of synchrony could account for observations of non-reciprocal relationships between simultaneously recorded PCs and the downstream target neurons they inhibit^45^. Our finding of very little synchrony in pairs of PCs suggests the potential importance of other neural circuit mechanisms to account for the potentially non-reciprocal relationships between PCs and their downstream target neurons. In general, neurons in the cerebellar nucleus receive excitatory inputs from mossy-fiber collaterals and other non-PC sources. Thus, their firing need not be driven solely by the inhibitory input from PCs and should reflect the balance of PC inhibition and excitation from non-PC inputs. It is easy to imagine that, in some circumstances, non-PC excitation might predominate over PC inhibition such that some PCs show increased firing at the same time as their target neurons. In the specific case of the oculomotor vermis, for instance, PC responses show a high degree of heterogeneity during saccades^27,37^. The oculomotor vermis is an example of regions that do not share consistent simple spike responses across PCs, in which case a reciprocal relationship between individual PCs and downstream neurons would not be expected. In the specific case of the cerebellar flocculus, PCs share similar direction preferences in simple spike encoding of pursuit eye movements^30,33^, and downstream FTNs show firing that is roughly the reciprocal of the PC population. Together, our data and these examples make us lean towards neural circuit mechanisms to create coordinated, rather than reciprocal, firing of cerebellar PCs and their target neurons in the cerebellar nuclei.

### Rigor in assessment of Purkinje cell synchrony

We were as rigorous as we could be in assessing PC synchrony as a potential neural code. Some PC synchrony is expected given the mean firing rates of simultaneously recorded neurons. For instance, two independent PCs with mean firing rates of 60 spikes/s should produce 3.6 synchronous (within 1 ms) spikes in a one second period. Our goal was to quantify whether PC synchrony was larger or smaller than would be expected given PC firing rates by estimating and subtracting the synchrony expected solely from changes in PC firing rates. In agreement with previous observations of synchrony between pairs of PCs in the cerebellum^18–27^, we indeed found that PCs synchronize slightly more than would be expected. Yet, the magnitude of PC-PC synchrony is relatively small, corresponding to a single extra synchronous spike between PC pairs over a period of 5-10 seconds. Similar to previous observations^19^, synchrony tended to be higher for nearby PCs, with significant heterogeneity in both the magnitude and timing of joint firing.

We also attempted to mitigate the challenges posed by spike-sorting in assaying synchrony from extra-cellular recordings. Spike-sorting is effectively a layer of statistical inference between the actual recordings and the assigned timing of the spikes of each neuron, and it could bias results either toward or away from synchrony. Temporal collisions between spikes, corresponding to millisecond-level synchrony between neurons, are particularly problematic for spike-sorters^35,46,47^, especially given the high degree of similarity between PC waveforms, the high intrinsic firing rates of PCs, and the observation that synchrony decreases with separation. Two features of our data make us believe that our quantification of synchrony is accurate. First, we designed a spike-sorter that would disambiguate temporal collisions between neurons and tested the sorter using data that closely mimicked our cerebellar recordings^35^. Second, in PC pairs where we observed millisecond synchrony, we also observed a pause in firing of approximately equal magnitude as both PCs enter their refractory periods. A post-synchrony pause is unexpected if the measured synchrony was due to the incorrect addition of a synchronous spike, as our sorting methodology does not enforce the presence of a neuron refractory period.

### Circuit mechanisms that might promote or counteract Purkinje cell synchrony

Given the number of potential circuit and cellular mechanisms that might promote PC synchrony, we were surprised to record so little. For instance, recordings from PCs separated by ~20 um in the mouse show strong synchrony even in the absence of synaptic input^19^, suggesting that ephaptic coupling may synchronize the simple spikes of nearby PCs. In addition, PC synchrony could be inherited from upstream granule cell synchrony that may, in turn, be a fundamental property of mossy fiber inputs to the cerebellum or derived from extensive innervation of granule cells by a single mossy fiber in the glomerulus^48^. The dramatic expansion of coding in the granule cells, coupled with ephaptic connections between adjacent PCs might tend to promote PC synchrony. Yet, we suggest that just as some circuit mechanisms within the cerebellar cortex could promote synchrony, others may directly counter synchrony. For instance, molecular layer interneurons (MLIs) receive granule cell input and inhibit PCs^49^, thus potentially suppressing the effects of synchronous granule cells inputs to PCs. Ephaptic coupling from some MLIs (basket cells) to PCs^50^ might allow rapid inhibition of PCs from synchronous granule cell inputs, an effect that could be amplified by gap junction synapses between MLIs^51–53^. A balanced set of circuit mechanisms that both promote and constrain synchrony may allow the cerebellum to maintain a consistent output code even when the statistics of its input vary widely.

### Applicability to other cerebellar regions

Our results do not rule out the possibility that synchrony is a mechanism of information transmission in other areas of the cerebellum. For example, two prior studies have suggested that PC synchrony is modulated during motor behavior^26,27^. However, the common finding of small magnitudes of synchrony in our data and past studies^18–27^ suggests a consistent level of PC synchrony across regions, tasks, and species. Where differences exist, they could result from the metrics used to assay time-dependent changes in synchrony. Here, we took care to ensure that our metric of synchrony was uncontaminated by changes in firing rate and we ran multiple control analyses to ensure that our results were consistent across metrics.

Finally, synchrony could contribute to cerebellar signaling under some circumstances. Timing codes may be particularly relevant when mossy fiber inputs are highly synchronized, such as during a very brief stimulus^54^. We also note that a well-timed spike in downstream neurons could be accomplished by coordinated pauses in PC firing^55–57^ rather than via millisecond-precision synchronous spikes. Yet, caveats aside, we suggest that for most real-life circumstances during behavior, Purkinje cells affect the firing of downstream neurons primarily through modulation of the rate of the simple spikes they use to communicate with neurons in the cerebellar nucleus.

## Methods

Three male rhesus monkeys (*Macaca mulatta*, 10-14 kg) served as experimental subjects. All experimental procedures were preapproved by the Institutional Animal Care and Use Committee at Duke University and followed the NIH *Guide for the Care and Use of Laboratory Animals* (1977). We also reanalyzed previously reported^31^ experiment data from FTNs in the vestibular nucleus (n=2 monkeys).

### General procedures

All monkeys underwent several separate surgical procedures that used sterile technique. During all surgeries, monkeys were deeply anesthetized with isoflurane. Monkeys received analgesics following each procedure until they had recovered. In an initial surgery, a head-holder was implanted to minimize motion of the monkey’s head during future neurophysiological recording sessions and to ensure that we could measure movements of the monkey’s eyes uncontaminated by head movement. In a second surgery, we sutured a coil of wire to the sclera of one eye^58^, allowing the recording of eye position and velocity with high spatial and temporal precision via the search coil technique^59^. Following these two surgeries, monkeys were trained to pursue a moving target in exchange for a liquid reward. Once monkeys demonstrated proficiency in tracking the dot (with minimal intervening saccadic eye movements), they underwent a final surgical procedure to implant a recording chamber and allow our electrodes to access the cerebellar floccular complex.

All experiments were performed in a dimly lit room while the monkey’s head was fixed 30 cm in front of a CRT monitor. Visual targets consisted of a black 0.5° diameter spot presented on a gray background. Motion of the dot on each trial was controlled by our laboratory’s custom “Maestro” software. During each trial, we recorded the voltages corresponding to the horizontal and vertical position of the monkey’s eyes, sampling at 1 kHz. Position signals were differentiated offline using the central difference method and subsequently low pass filtered using a non-causal 2^nd^ order Butterworth filter to produce estimates of the monkey’s eye velocity on each trial (cut-off frequency of 30 Hz). We used an automated procedure to identify saccades using a combination of eye velocity (20 deg/s) and acceleration thresholds (1,250 deg/s/s). Periods from 10 ms before to 10 ms after exceeding the joint velocity-acceleration thresholds were treated as missing data in all subsequent analyses.

### Cerebellar flocculus recording procedures

Each day, we acutely inserted either tungsten micro-electrodes or custom-designed 16-channel Plexon S-probes into the cerebellar flocculus. S-probes were 185 microns in diameter and featured 7.5-micron diameter tungsten contacts arranged in two columns of eight contacts with 50 micron spacing between adjacent rows and columns. We identified the floccular complex by its strong response to smooth pursuit eye movements as well as the presence of intermittent Purkinje cell complex spikes. Continuous wideband voltage measurements from all channels were recorded at 40 kHz using Plexon hardware. To ensure that wideband data was not contaminated by the electrical field produced by the eye coil drivers, we used a hardware based 4-pole low-pass Butterworth filter with a 6 kHz cut-off frequency prior to digitization by the Plexon system.

After arriving in the cerebellar flocculus and isolating one or more Purkinje cells, we allowed the electrode to rest for a minimum of 30 minutes (up to several hours). The waiting time ensured that any recorded units were maximally stabile for the duration of the recording. Any additional drift of the neural unit during the recording was corrected during spike-sorting (see spike-sorting procedures, below). The close spacing of the multi-contact probe’s contacts and our spike-sorting strategy ensured that we were able to track units should they move across contacts during the recording session.

### Spike-sorting and quality control metrics

After each recording session, we assigned recorded extra-cellular spikes to individual neural units using the semi-automated “Full Binary Pursuit” (FBP) spike-sorter^35^. As we were especially interested in quantifying the magnitude of synchrony between pairs of simultaneously recorded PCs, we chose the FBP sorter due to its superior performance in deconflating spike collisions on multi-contact electrodes. Briefly, we used a zero-phase FIR bandpass filter between 300 Hz and 8 kHz to isolate the action potentials from lower frequency signals. Temporally isolated actional potentials were identified on each channel by clipping the voltage 0.3 ms before to 0.7 ms after a peak voltage deflection. These isolated action potentials were then clustered using a modification of the iso-cut algorithm^60^ to identify the waveform signatures (templates) of individual neurons across channels. To identify the spike times of individual units in the face of temporally and spatially overlapping spikes, the voltage timeseries at each timepoint was modeled as the potential sum of one or more of the identified neuron templates across channels (“binary pursuit”)^35,61^.

After automated sorting, we manually curated the output of the sorting algorithm by removing any neurons with significant ISI violations (operationally defined as the fraction of a neuron’s spikes that occurred within 1 ms of each other). In our dataset, identified PCs had very few ISI violations: 0.33 ± 0.56% (mean ± std.), indicating the ability of the sorter to appropriately classify the spikes of isolated Purkinje cells. We additionally removed any units with a low signal-to-noise (SNR) ratio. We defined the SNR as the voltage range spanned by the neuron’s template divided by the standard deviation of the background noise on the neuron’s primary channel. As the majority of neurons exhibit substantial voltage deflections on multiple channels, the definition of SNR we used represents a lower bound on our estimate of the isolation quality of a single unit. We also note that use of the conservative definition of SNR ensures that substantial amounts of drift, in which the neuron’s primary channel might change across the recording session, always resulted in a reduction in our estimate of the units SNR. Across our dataset, PCs had a SNR of 9.02 ± 3.99 (mean ± SD). For analyses that used estimates of a neuron’s continuous firing rate, we convolved a causal double-exponential filter with the recorded neuron’s spike train (*τ_rise_* = 0.1 ms, *τ_decay_* = 50 ms).

### Identification of Purkinje cells

Purkinje cells receive a single climbing fiber input from the inferior olive. Post-synaptic climbing fiber responses drive PC complex spikes, the occurrence of which results in a stereotypical simple spike pause for 10s of milliseconds. Our basis for labeling a floccular neuron as “known PC” was based on the presence of well-defined pause in the co-recorded simple spikes following an identified complex spike. Our data include *n=110* well-isolated known PCs, including *n=32* simultaneously recorded pairs. We also recorded many neurons that satisfied all criteria as a PC except that we were not able to isolate the complex spike simultaneously and therefore assay the presence of a post-CS pause in simple-spike firing. Our recordings included *n=49* pairs that included a simultaneously recorded putative PC with a known PC as well as *n=36* additional pairs where both members of the pair were putative PCs. The combined sample of *n=85* “putative PC pairs” formed a non-overlapping set where we could explicitly test whether these pairs featured synchrony properties that were similar to those in our *n=32* pairs of known PCs.

### Behavioral procedures

We measured the monkey’s eye kinematics and neuronal responses during discrete trials of smooth target motion. At the start of each trial, the monkey fixated the stationary visual target for a random interval between 400 and 800 ms. Then, the target instantaneously jumped backwards by 3-5 degrees and began moving in the opposite direction at a constant speed for 650 ms (using the “step-ramp” paradigm of Rashbass^62^ to minimize catch-up saccades during pursuit initiation). At the end of each trial, the monkey fixated the stationary target eccentrically for an additional 200 ms. In exchange for appropriate tracking of the target as well as fixation at the beginning and end of each experimental trial (within an invisible bounding box extending ±3 degrees from the target), monkeys received a small liquid reward. If the monkey broke fixation during the fixation interval or failed to adequately track the target, the trial immediately aborted and the monkey was not rewarded. Aborted trials were not included in the data analysis.

### Identification of temporal synchrony between simultaneously recorded Purkinje cells

Our goal was to identify, as a function of time during repetitions of smooth pursuit tracking, the presence of any unexpected temporal structure between pairs of Purkinje cells beyond what would be expected due to their time varying firing rates. Our ideal metric for quantifying unexpected temporal relationships between PCs would satisfy three criteria. First, the metric should be model-free, not requiring an assumption about the exact timescale of temporal interactions between simultaneously recorded neurons. Second, the same procedure should be capable of identifying unexpected temporal relationships between neurons from both continuous spiking data as well as measured across repeated presentations of the same behavioral stimulus. Most importantly, the metric should not be biased by changes in either the mean (background) firing rates of the two neurons or by the temporally slower changes in firing rates due to stimuli/behavior. To satisfy these three criteria, we devised a procedure for estimating the temporal relationships between simultaneously recorded Purkinje cells while accounting for local changes in firing rate. Our procedure relies on two primary assumptions: (1) there is sufficient independent noise in the timing of Purkinje cell spikes so that repeated presentations of the same behavioral stimulus result in temporally-jittered spiking across trials and (2) temporal coordination between cells should manifest as consistent deviations in the spike timing of one cell relative to another. Our assumption about sufficient independent spike timing variability ensures that temporal ordering effects between two Purkinje cells are not driven by the stimulus or behavior.

Our procedure to identify temporal coordination between simultaneously recorded neurons relies on comparison of the raw synchrony with the probability of observing a spike in millisecond bins under the null hypothesis that spike timing is dependent only on the local firing rate of the neuron under study.

To compute the null-hypothesis, we consider a set of three consecutive spikes in PC1 with times *T_i-1_, T_i_*, and *T_i+1_*. The spike at *T_i_* could be anywhere from backwards by up to half of the [*T_i_, T_i+1_* interval or forwards by half of the subsequent [*T_i_, T_i+1_* interval without altering the average firing rate over the *T_i-1_* to *T_i+1_* interval. We took advantage of the stability of average firing rate over the *T_i-1_* to *T_i+1_* interval to jitter the spikes by distributing their probability uniformly over the interval between halfway from the previous spike and halfway to the subsequent spike and thereby created 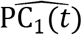, the expected probability of observing a spike in PC1 across time, dependent solely on its local firing rate:

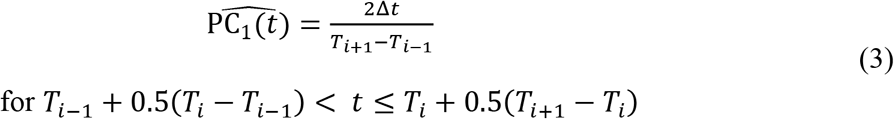

In Equation (3), *T_i_* is the timing of PC1’s *i^th^* spike and Δ*t* is the binwidth of 1 ms. While Equation (3) assumes a uniform probability of spiking between adjacent interspike intervals, other distributions of spike probability could be used (e.g., exponential, gamma, or empirically derived interspike interval distributions). As our conclusions about PC synchrony are not strongly dependent on the choice of interspike interval distribution, we chose to assume a uniform distribution of spike timings. 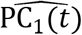 thus quantifies the probability of a spike occurring in each millisecond bin under the null hypothesis that spike timing is dictated solely by the local firing rate.

To quantify the raw synchrony as a function of time during pursuit, we computed the intersection of the two PC spike trains across time (Pr[PC_1_(*t*) ⋂ PC_2_(*t*)]). We then averaged the intersection of the PC spike trains across replicates of pursuit trials with the same pursuit stimulus to obtain the mean probability of observing synchronous spikes at each millisecond. To assay if the observed probability of synchronous spikes deviated from what would be expected based on the PC’s time-varying firing rates, we computed 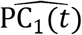 and 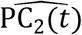 for each smooth pursuit trial according to Equation (3). The expected probability of synchrony due to the independent variation of firing rates can then be computed as the product of the two null hypothesized probability timeseries.

### Calculation of cross-correlograms

For each pair of simultaneously recorded PCs, we generated a cross-correlogram (CCG) between the two neurons without regard for time during a behavior using Equation 1. Our goal was to identify whether the presence of a spike in the trigger neuron (PC2) influenced the timing of PC1’s spikes. Yet, the CCG contains both temporal information related rapid temporal coordination between the two neurons and lower frequency relationships due to co-modulation of firing patterns. To specifically identify the more rapid timescales corresponding to temporal coordination between PC pairs, we removed rate-based effects. First, we generated the probability of observing a spike for PC1 in each millisecond bin under the null hypothesis that spiking timing was dependent solely on the local firing rate via Equation (3). Then, we computed a rate-corrected CCG by explicitly removing the null hypothesized probability of spiking triggered on the timing of PC2’s spikes:

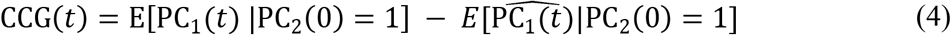

Any spiking of PC2 that consistently affects the timing of spikes for PC1 would violate the null hypothesis and create a deviation in *CCG(t*). This method has the advantage that it allows us to generate confidence intervals for each bin by asking whether the number of spikes in each bin obeys a binomial distribution with a mean probability defined from the null distribution at each timepoint.

Given a pair of PCs, the choice of the trigger neuron when generating the cross correlograms is ambiguous. To resolve the ambiguity, we generated two CCGs for each PC pair: one CCG where a randomly chosen neuron in the pair was the trigger neuron and a second CCG where the opposite neuron was considered the trigger. For each of these CCGs, we computed the mean of the CCG for all points *t* > 0 ms and chose the CCG with the smaller mean firing rate for *t* > 0 ms. Thereby, we ensured that for PCs that exhibited synchronous spiking at *t* = 0 ms, any decrease in the coincident firing of the two PCs would consistently occur for positive values of t. We note that alternative methods exist to choose the trigger neuron for generation of CCGs. For instance, one could either randomly choose the trigger neuron or compute a CCG with each PC in the pair serving as the trigger neuron and subsequently average the result. Either of these choices of trigger neuron would result in a more symmetric population CCG than our preferred method but would also obscure the clear pause after synchronous firing that is highlighted by our method. As we were primarily concerned with PC synchrony on the order of 1 ms, we ensured that our method of selecting the trigger neuron did not bias our estimate of excess synchrony measured at *t*=0 ms compared to a population of bootstrapped CCGs in which we randomly choose the trigger neuron.

### Alternative metrics for evaluating paired synchrony

Multiple methods have been proposed to evaluate temporal coordination and synchrony between simultaneously recorded pairs of neurons. Crucial for our endeavor was evaluating the magnitude of unexpected synchrony. Yet, many alternative metrics for assaying synchrony such as the correlation coefficient or pair-wise covariance are biased by changes in neuron firing rates (for a review, see ref. ^63^). One solution that accounts for biases induced by changes in firing rates is the “jitter-correction” method^64,65^. Most jitter-corrections specify the time scale of interest by setting the bin-width for jittering spike times. In contrast, our method described in Equation (3), does not require an explicit assumption about the time scale of interest. Instead, it explicitly tests whether the timing of PC1’s spikes are preferentially biased by the spiking of PC2.

A second class of methods for assaying synchrony relies on multiple trial replicates, with the goal of removing the contributions of mean changes in firing rate measured across trial repeats from the joint probability of firing. Such trial-based synchrony metrics include the joint peristimulus time histogram^66^ (JPSTH), the synchrony index (SI)^19,27^, and the shift-predictor. The shift-predictor has a long been a metric for assaying synchrony across trials. Here, we used the shift-predictor as a control analysis to ensure that the use of our preferred method of hypothesis testing via Equation (3) was not missing synchrony that might otherwise exist.

Several previous reports have favored the synchrony index as a metric for assaying the magnitude of PC-PC synchrony during behavior^19,27^. While previous results have documented several short-comings of the SI metric (see, for instance, Equation 6 in ref. ^66^), we sought to explicitly test whether our data were consistent with past studies that relied on this metric. The synchrony index calculates the probability of observing simultaneous spikes from a pair of neurons, relative to the product of their independent probabilities. That is,

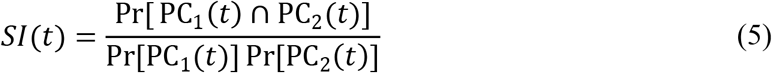

Here, Pr[PC_1_(*t*) ⋂ PC_2_(*t*)] represents the probability of observing a set of intersecting spikes between two PCs (PC1 and PC2) as a function of time across a set of trials. The probability of intersecting spikes is then normalized by the product of the timeseries of independent spiking probabilities. The synchrony index is exactly 1.0 when the two units fire independently. Values of the synchrony index that exceed 1.0 have been interpreted as synchrony that exceeds what would be expected by independent firing.

However, the SI conflates the measurement of two quantities: the firing rates of the individual units across time and the covariance of these units. When assaying spike timing synchrony as a function of trials, we really want to assess changes in the covariance of PC spike trains independent of changes in rate. The derivation below explains.

As PC_1_(*t*) and PC_2_(*t*) are binary spike trains, we can rewrite Pr[PC_1_(*t*) ⋂ PC_2_(*t*)] as the expected value of the product of these two timeseries: *E*[PC_1_(*t*)PC_2_(*t*)]. Similarly, Pr[PC_1_(*t*)] = *E*[PC_2_(*t*)] and Pr[PC_2_(*t*)] = *E*[PC_2_(*t*)]. The covariance between PC1 and PC2 is defined as:

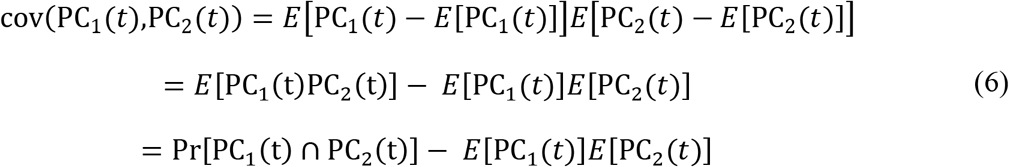

Thus, Pr[PC_1_(*t*) ⋂ PC_2_(*t*)] = cov(PC_1_(*t*), PC_2_(*t*)) + *E*[PC_1_(*t*)]*E*[PC_2_(*t*)] and we can rewrite synchrony index as:

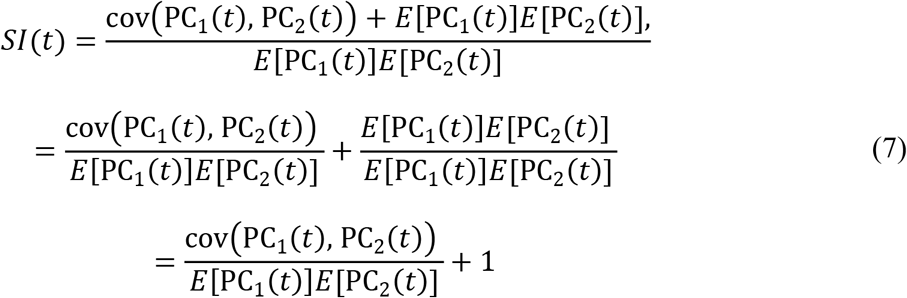

Equation (7) demonstrates that the synchrony index metric quantifies the covariance between the two simultaneously recorded PCs divided by the product of their firing rates. The synchrony index will increase if the product of the firing rates decreases, even if the covariance between the two units is unchanged. Our analysis in Supplementary Figure 1 shows how changes in the firing rates of simultaneously recorded units can produce changes in the measured synchrony index in the absence of any temporal modulation in the covariance of the two neurons’ spike trains. The conflation of covariance and firing rate might explain a previous report of an increase in the synchrony index at points in a movement when population firing rates in the oculomotor vermis decrease^27^.

### Simulations of independent and covarying Purkinje cell populations

To quantify the effect of spike timing synchrony on downstream neurons in the cerebellar nuclei, we constructed two simulated populations of PCs. Each population consisted of *n=40* simulated PC spike trains. The sum of these spike trains was taken as the input to a hypothetical downstream neuron in the cerebellar nucleus. We randomly assigned the mean firing rate of each simulated PC by selecting a mean firing rate from one of our *n=110* known PCs. For each neuron in an “independent” population, we generated a random Poisson distributed spike train with a temporal resolution of 1 millisecond. For each simulated PC-PC pair in a “non-zero covariance” population, we generated spike trains by randomly assigning a covariance value from our empirical distribution measured from each pair of PCs in our recorded population. We found this empirical distribution of covariance values by using the value of the rate-corrected CCGs at *t=0* ms, scaled by the probability of observing a spike in the trigger neuron. The scaling of the rate-corrected CCG resulted in a rate-corrected pair-wise covariance that accounted for fluctuations in the mean firing rates of the two PCs. Randomly choosing pairwise covariances from our recorded population may not always produce a valid covariance matrix across the complete population. If the resulting covariance matrix was not positive semi-definite, we found the closest covariance matrix using the Higham iterative correction^67^. Then, we computed Poisson distributed PC spike trains under the corrected covariance matrix by pulling from a multidimensional Gaussian distribution using the procedure found in Macke et al.^68^. After generating spike trains for both the independent and non-zero covariance populations, we measured the mean number of spikes that would arrive at a downstream neuron contingent on the occurrence of a spike in any of the simulated PCs.

Previous experiments using dynamic clamp^13^ quantified the effect of PC synchrony on entrainment of downstream neurons by exactly simulating the spiking of a population of PCs as inputs to a recorded cerebellar nucleus neuron. The experiments controlled the fraction of the simulated PC population that was “fully synchronous” and asked how the level of synchrony affected downstream firing rates. To aid in comparison with these past results, we asked what fraction of our PC population would need to be fully synchronous to match the mean number of spikes that arrive at the downstream population given that one of the simulated PCs fired. If all PCs are independent, then one spike in one PC would be accompanied, on average, by 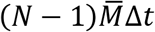 additional synchronous spikes from the point of view of a downstream neuron. Here, *N* is the number of neurons in the PC population (*N*=40), 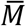 is the mean firing rate of the PCs in spikes/s, and Δ*t* is the temporal resolution (1 ms). If all PCs are fully synchronous, then the firing of one PC completely predicts the firing of the rest of the population, resulting in (*N* – 1) spikes arriving at the downstream neuron simultaneously. We solved this model for all values of synchrony. Let *x* be the fraction of the population that is fully synchronous, ranging from 0 (fully independent) to 1 (completely synchronous). Then, the firing of a single PC in the population predicts the arrival of additional spikes at the downstream neuron according to Equation 8:

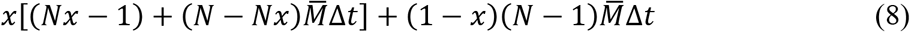

### Statistical analysis

Unless otherwise noted, all statistical tests were two-tailed and used a significance level of p < 0.05. Statistical analyses were performed using the HypothesisTests package in Julia.

## Acknowledgements

This research was supported by NIH grants R01-NS112917 (SGL) and K99-EY030528 (DJH). We thank Stefanie Tokiyama and Bonnie Bowell for technical assistance, including care of the monkeys.

## Data availability

Purkinje cell spike trains, associated behavior data, and summary data plotted in all figures are available via the Open Science Framework repository (osf.io/wjg32).

---

**Supplementary Figure 1.**
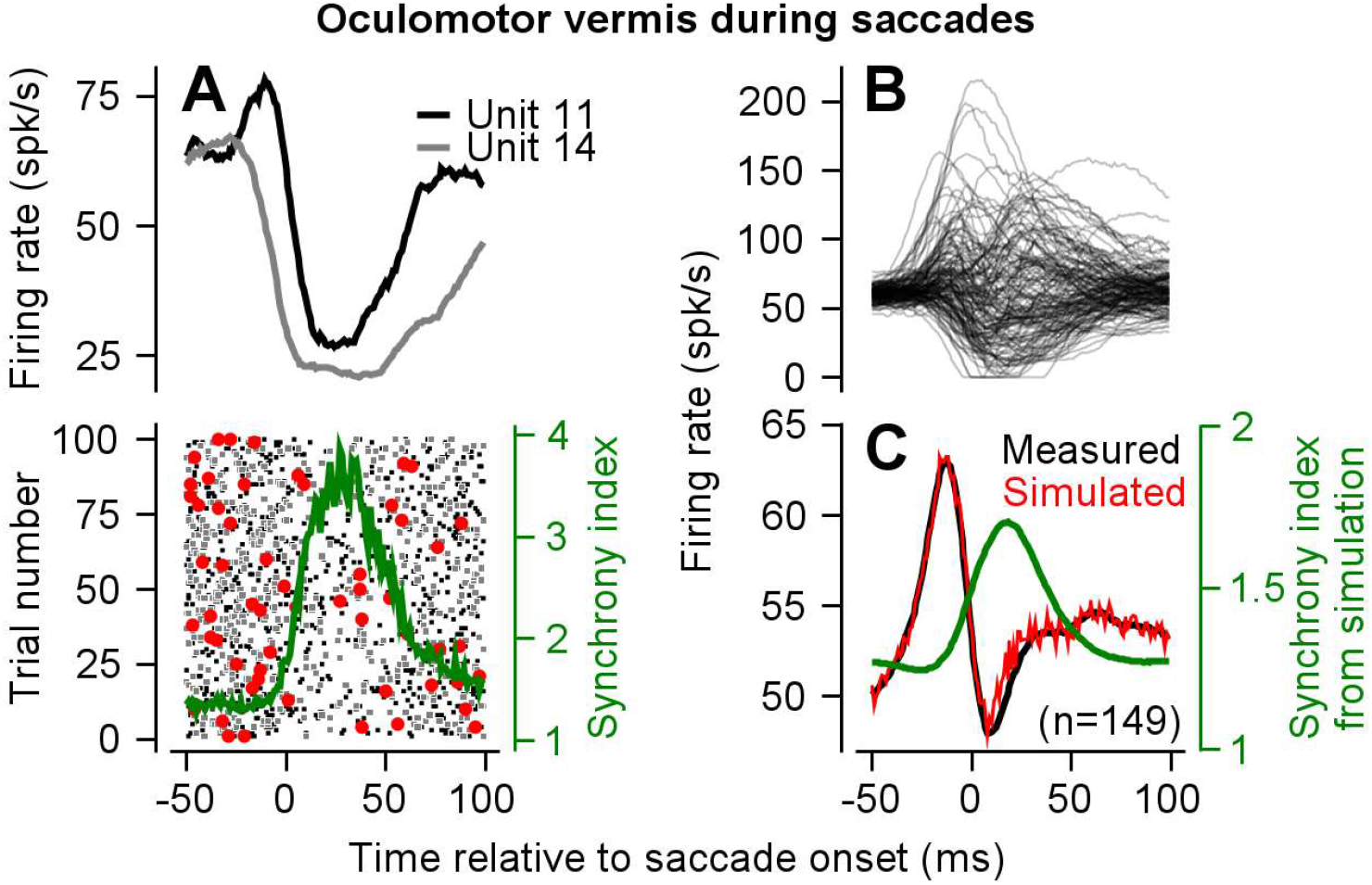
The synchrony index conflates spike timing covariance and mean firing rate. **A.** Top graph shows example firing rate traces (top) for two Purkinje cells recorded from the oculomotor vermis of marmosets (*callitrix jacchus*) during execution of a saccadic eye movement, taken from publicly available data from reference ^27^. Mean firing rate from that report were offset by 50 spikes/s to match baseline rates typical of PCs in the cerebellum. Bottom plot shows simulated raster plots across n=100 trials for both units, each following the mean firing rate plotted above. Spike times were drawn such that the covariance between the two simulated neurons was constant as a function of time (0.0015). Spikes that occur in the two simulated spike trains within the same millisecond window are shown in red. Green curve shows the synchrony index computed for these two units. **B.** Firing rate responses of *n=149* PCs from marmosets during saccadic eye movements, each with an assumed 50 spikes/s baseline rate, again taken from reference 41. **C.** Mean population response averaged across the neurons shown in (B). Red curve shows simulated mean response with a constant covariance between units (0.0015). Green curve shows the synchrony index measured across the simulated population. Results in (C) do not depend qualitatively on the mean or variance of the assumed baseline firing rate.

## References

1. Adrian, E. D. & Matthews, R. The action of light on the eye. J. Physiol. 65, 273–298 (1928).

2. Bagnall, M. W., McElvain, L. E., Faulstich, M. & Lac, S. du. Frequency-Independent Synaptic Transmission Supports a Linear Vestibular Behavior. Neuron 60, 343–352 (2008).

3. Turecek, J., Jackman, S. L. & Regehr, W. G. Synaptic Specializations Support Frequency-Independent Purkinje Cell Output from the Cerebellar Cortex. Cell Rep. 17, 3256–3268 (2016).

4. Gollisch, T. & Meister, M. Rapid Neural Coding in the Retina with Relative Spike Latencies. Science 319, 1108–1111 (2008).

5. Junek, S., Kludt, E., Wolf, F. & Schild, D. Olfactory Coding with Patterns of Response Latencies. Neuron 67, 872–884 (2010).

6. Hahnloser, R. H. R., Kozhevnikov, A. A. & Fee, M. S. An ultra-sparse code underliesthe generation of neural sequences in a songbird. Nature 419, 65–70 (2002).

7. Chase, S. M. & Young, E. D. Spike-Timing Codes Enhance the Representation of Multiple Simultaneous Sound-Localization Cues in the Inferior Colliculus. J. Neurosci. 26, 3889–3898 (2006).

8. Chase, S. M. & Young, E. D. Cues for Sound Localization Are Encoded in Multiple Aspects of Spike Trains in the Inferior Colliculus. J. Neurophysiol. 99, 1672–1682 (2008).

9. Sober, S. J., Sponberg, S., Nemenman, I. & Ting, L. H. Millisecond Spike Timing Codes for Motor Control. Trends Neurosci. 41, 644–648 (2018).

10. Diedrichsen, J., King, M., Hernandez-Castillo, C., Sereno, M. & Ivry, R. B. Universal Transform or Multiple Functionality? Understanding the Contribution of the Human Cerebellum across Task Domains. Neuron 102, 918–928 (2019).

11. Schmahmann, J. D. From movement to thought: Anatomic substrates of the cerebellar contribution to cognitive processing. Hum. Brain Mapp. 4, 174–198 (1996).

12. Eccles, J. C., Llinás, R. & Sasaki, K. The excitatory synaptic action of climbing fibres on the Purkinje cells of the cerebellum. J. Physiol. 182, 268–296 (1966).

13. Person, A. L. & Raman, I. M. Purkinje neuron synchrony elicits time-locked spiking in the cerebellar nuclei. Nature 481, 502–505 (2012).

14. Raman, I. M., Gustafson, A. E. & Padgett, D. Ionic currents and spontaneous firing in neurons isolated from the cerebellar nuclei. J. Neurosci. Off. J. Soc. Neurosci. 20, 9004–9016 (2000).

15. Jahnsen, H. Electrophysiological characteristics of neurones in the guinea-pig deep cerebellar nuclei in vitro. J. Physiol. 372, 129–147 (1986).

16. Wu, Y. & Raman, I. M. Facilitation of mossy fibre-driven spiking in the cerebellar nuclei by the synchrony of inhibition. J. Physiol. 595, 5245–5264 (2017).

17. Hoebeek, F. E., Witter, L., Ruigrok, T. J. H. & Zeeuw, C. I. D. Differential olivo-cerebellar cortical control of rebound activity in the cerebellar nuclei. Proc. Natl. Acad. Sci. 107, 8410–8415 (2010).

18. de Solages, C. et al. High-Frequency Organization and Synchrony of Activity in the Purkinje Cell Layer of the Cerebellum. Neuron 58, 775–788 (2008).

19. Han, K.-S.et al. Ephaptic Coupling Promotes Synchronous Firing of Cerebellar Purkinje Cells. Neuron 100, 564–578.e3 (2018).

20. Bell, C. C. & Grimm, R. J. Discharge properties of Purkinje cells recorded on single and double microelectrodes. J. Neurophysiol. 32, 1044–1055 (1969).

21. Bell, C. C. & Kawasaki, T. Relations among climbing fiber responses of nearby Purkinje Cells. J. Neurophysiol. 35, 155–169 (1972).

22. MacKay, W. A. & Murphy, J. T. Integrative versus delay line characteristics of cerebellar cortex. Can. J. Neurol. Sci. J. Can. Sci. Neurol. 3, 85–97 (1976).

23. Ebner, T. J. & Bloedel, J. R. Correlation between activity of Purkinje cells and its modification by natural peripheral stimuli. J. Neurophysiol. 45, 948–961 (1981).

24. De Zeeuw, C. I., Koekkoek, S. k. e., Wylie, D. r. w. & Simpson, J. I. Association Between Dendritic Lamellar Bodies and Complex Spike Synchrony in the Olivocerebellar System. J. Neurophysiol. 77, 1747–1758 (1997).

25. Wise, A. K., Cerminara, N. L., Marple-Horvat, D. E. & Apps, R. Mechanisms of synchronous activity in cerebellar Purkinje cells. J. Physiol. 588, 2373–2390 (2010).

26. Heck, D. H., Thach, W. T. & Keating, J. G.On-beam synchrony in the cerebellum as the mechanism for the timing and coordination of movement. Proc. Natl. Acad. Sci. 104, 7658–7663 (2007).

27. Sedaghat-Nejad, E., Pi, J. S., Hage, P., Fakharian, M. A. & Shadmehr, R. Synchronous spiking of cerebellar Purkinje cells during control of movements. Proc. Natl. Acad. Sci. 119, e2118954119 (2022).

28. Rambold, H., Churchland, A., Selig, Y., Jasmin, L. & Lisberger, S. G. Partial ablations of the flocculus and ventral paraflocculus in monkeys cause linked deficits in smooth pursuit eye movements and adaptive modification of the VOR. J. Neurophysiol. 87, 912–924 (2002).

29. Lisberger, S. G. & Fuchs, A. F. Role of primate flocculus during rapid behavioral modification of vestibuloocular reflex. I. Purkinje cell activity during visually guided horizontal smooth-pursuit eye movements and passive head rotation. J. Neurophysiol. 41, 733–763 (1978).

30. Medina, J. F. & Lisberger, S. G. Variation, Signal, and Noise in Cerebellar Sensory–Motor Processing for Smooth-Pursuit Eye Movements. J. Neurosci. 27, 6832–6842 (2007).

31. Joshua, M., Medina, J. F. & Lisberger, S. G. Diversity of Neural Responses in the Brainstem during Smooth Pursuit Eye Movements Constrains the Circuit Mechanisms of Neural Integration. J. Neurosci. 33, 6633–6647 (2013).

32. Stone, L. S. & Lisberger, S. G. Visual responses of Purkinje cells in the cerebellar flocculus during smooth-pursuit eye movements in monkeys. I. Simple spikes. J. Neurophysiol. 63, 1241–1261 (1990).

33. Krauzlis, R. J. & Lisberger, S. G. Simple spike responses of gaze velocity Purkinje cells in the floccular lobe of the monkey during the onset and offset of pursuit eye movements. J. Neurophysiol. 72, 2045–2050 (1994).

34. Lisberger, S. G., Pavelko, T. A. & Broussard, D. M. Responses during eye movements of brain stem neurons that receive monosynaptic inhibition from the flocculus and ventral paraflocculus in monkeys. J. Neurophysiol. 72, 909–927 (1994).

35. Hall, N. J., Herzfeld, D. J. & Lisberger, S. G. Evaluation and resolution of many challenges of neural spike sorting: a new sorter. J. Neurophysiol. 126, 2065–2090 (2021).

36. Shadmehr, R. Population coding in the cerebellum: a machine learning perspective. J. Neurophysiol. 124, 2022–2051 (2020).

37. Herzfeld, D. J., Kojima, Y., Soetedjo, R. & Shadmehr, R. Encoding of action by the Purkinje cells of the cerebellum. Nature 526, 439–442 (2015).

38. Herzfeld, D. J., Kojima, Y., Soetedjo, R. & Shadmehr, R. Encoding of error and learning to correct that error by the Purkinje cells of the cerebellum. Nat. Neurosci. 21, 736–743 (2018).

39. Sugihara, I., Wu, H.-S. & Shinoda, Y. The Entire Trajectories of Single Olivocerebellar Axons in the Cerebellar Cortex and their Contribution to Cerebellar Compartmentalization. J. Neurosci. 21, 7715–7723 (2001).

40. De Zeeuw, C. I.et al. Spatiotemporal firing patterns in the cerebellum. Nat. Rev. Neurosci. 12, 327–344 (2011).

41. Ozden, I., Sullivan, M. R., Lee, H. M. & Wang, S. S.-H. Reliable Coding Emerges from Coactivation of Climbing Fibers in Microbands of Cerebellar Purkinje Neurons. J. Neurosci. 29, 10463–10473 (2009).

42. Zee, D. S., Yamazaki, A., Butler, P. H. & Gucer, G. Effects of ablation of flocculus and paraflocculus of eye movements in primate. J. Neurophysiol. 46, 878–899 (1981).

43. Medina, J. F. & Lisberger, S. G. Encoding and decoding of learned smooth-pursuit eye movements in the floccular complex of the monkey cerebellum. J. Neurophysiol. 102, 2039–2054 (2009).

44. Voogd, J., Pardoe, J., Ruigrok, T. J. H. & Apps, R. The Distribution of Climbing and Mossy Fiber Collateral Branches from the Copula Pyramidis and the Paramedian Lobule: Congruence of Climbing Fiber Cortical Zones and the Pattern of Zebrin Banding within the Rat Cerebellum. J. Neurosci. 23, 4645–4656 (2003).

45. McDevitt, C. J., Ebner, T. J. & Bloedel, J. R. Relationships between simultaneously recorded Purkinje cells and nuclear neurons. Brain Res. 425, 1–13 (1987).

46. Garcia, S., Buccino, A. P. & Yger, P. How Do Spike Collisions Affect Spike Sorting Performance? eNeuro 9, (2022).

47. Bar-Gad, I., Ritov, Y., Vaadia, E. & Bergman, H. Failure in identification of overlapping spikes from multiple neuron activity causes artificial correlations. J. Neurosci. Methods 107, 1–13 (2001).

48. Nguyen, T. M.et al. Structured cerebellar connectivity supports resilient pattern separation. Nature 1–7 (2022) doi:10.1038/s41586-022-05471-w.

49. Eccles, J. C. The Cerebellum as a Neuronal Machine. (Springer-Verlag, 1967).

50. Blot, A. & Barbour, B. Ultra-rapid axon-axon ephaptic inhibition of cerebellar Purkinje cells by the pinceau. Nat. Neurosci. 17, 289–295 (2014).

51. Kozareva, V. et al. A transcriptomic atlas of mouse cerebellar cortex comprehensively defines cell types. Nature 598, 214–219 (2021).

52. Mann-Metzer, P. & Yarom, Y. Electrotonic coupling interacts with intrinsic properties to generate synchronized activity in cerebellar networks of inhibitory interneurons. J. Neurosci. Off. J. Soc. Neurosci. 19, 3298–3306 (1999).

53. Mann-Metzerand, P. & Yarom, Y. Electrotonic coupling synchronizes interneuron activity in the cerebellar cortex. in Progress in Brain Research vol. 124 115–122 (Elsevier, 2000).

54. Brown, S. T. & Raman, I. M. Sensorimotor Integration and Amplification of Reflexive Whisking by Well-Timed Spiking in the Cerebellar Corticonuclear Circuit. Neuron 99, 564–575.e2 (2018).

55. Gauck, V. & Jaeger, D. The Control of Rate and Timing of Spikes in the Deep Cerebellar Nuclei by Inhibition. J. Neurosci. 20, 3006–3016 (2000).

56. Steuber, V. & Jaeger, D. Modeling the generation of output by the cerebellar nuclei. Neural Netw. 47, 112–119 (2013).

57. Steuber, V. et al. Cerebellar LTD and Pattern Recognition by Purkinje Cells. Neuron 54, 121–136 (2007).

58. Ramachandran, R. & Lisberger, S. G. Normal Performance and Expression of Learning in the Vestibulo-Ocular Reflex (VOR) at High Frequencies. J. Neurophysiol. 93, 2028–2038 (2005).

59. Robinson, D. A. A Method of Measuring Eye Movemnent Using a Scieral Search Coil in a Magnetic Field. IEEE Trans. Bio-Med. Electron. 10, 137–145 (1963).

60. Chung, J. E.et al. A Fully Automated Approach to Spike Sorting. Neuron 95, 1381–1394.e6 (2017).

61. Pillow, J. W., Shlens, J., Chichilnisky, E. J. & Simoncelli, E. P. A Model-Based Spike Sorting Algorithm for Removing Correlation Artifacts in Multi-Neuron Recordings. PLOS ONE 8, e62123 (2013).

62. Rashbass, C. The relationship between saccadic and smooth tracking eye movements. J. Physiol. 159, 326–338 (1961).

63. Cutts, C. S. & Eglen, S. J. Detecting Pairwise Correlations in Spike Trains: An Objective Comparison of Methods and Application to the Study of Retinal Waves. J. Neurosci. 34, 14288–14303 (2014).

64. Amarasingham, A., Harrison, M. T., Hatsopoulos, N. G. & Geman, S. Conditional modeling and the jitter method of spike resampling. J. Neurophysiol. 107, 517–531 (2011).

65. Smith, M. A. & Kohn, A. Spatial and Temporal Scales of Neuronal Correlation in Primary Visual Cortex. J. Neurosci. 28, 12591–12603 (2008).

66. Aertsen, A. M., Gerstein, G. L., Habib, M. K. & Palm, G. Dynamics of neuronal firing correlation: modulation of ‘effective connectivity’. J. Neurophysiol. 61, 900–917 (1989).

67. Higham, N. J. Computing the nearest correlation matrix—a problem from finance. IMA J. Numer. Anal. 22, 329–343 (2002).

68. Macke, J. H., Berens, P., Ecker, A. S., Tolias, A. S. & Bethge, M. Generating Spike Trains with Specified Correlation Coefficients. Neural Comput. 21, 397–423 (2009).

